# Selective integration during sequential sampling in posterior neural signals

**DOI:** 10.1101/642371

**Authors:** Fabrice Luyckx, Bernhard Spitzer, Annabelle Blangero, Konstantinos Tsetsos, Christopher Summerfield

## Abstract

Decisions are typically made after integrating information about multiple attributes of alternatives in a choice set. The computational mechanisms by which this integration occurs have been a focus of extensive research in humans and other animals. Where observers are obliged to consider attributes in turn, a framework known as “selective integration” can capture salient biases in human choices. The model proposes that successive attributes compete for processing resources and integration is biased towards the alternative with the locally preferred attribute. Quantitative analysis shows that this model, although it discards choice-relevant information, is optimal when the observers’ decisions are corrupted by noise that occurs beyond the sensory stage. Here, we used scalp electroencephalographic (EEG) recordings to test a neural prediction of the model: that locally preferred attributes should be encoded with higher gain in neural signals over posterior cortex. Over two sessions, human observers (of either sex) judged which of two simultaneous streams of bars had the higher (or lower) average height. The selective integration model fit the data better than a rival model without bias. Single-trial analysis showed that neural signals contralateral to the preferred attribute covaried more steeply with the decision information conferred by locally preferred attributes. These findings provide neural evidence in support of selective integration, complementing existing behavioural work.

**Significance Statement:** We often make choices about stimuli with multiple attributes, such as when deciding which car to buy on the basis of price, performance and fuel economy. A model of the choice process, known as selective integration, proposes that rather than taking all of the decision-relevant information equally into account when making choices, we discard or overlook a portion of it. Although information is discarded, this strategy can lead to better decisions when memory is limited. Here, we test and confirm predictions of the model about the brain signals that occur when different stimulus attributes of stimulus are being evaluated. Our work provides the first neural support for the selective integration model.

## INTRODUCTION

Biological brains evolved to be both precise and efficient. Responding accurately to external stimulation requires noisy sensory signals *x* to be transduced such that internal estimates 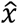 are as close as possible to their generative counterparts. Sequential sampling and integration allow the precision of sensory estimates to grow with the number of independent observations obtained (Wald and Wolfowitz, 1949), and neural signals in multiple brain regions implement a basic form of memory that allows optimal estimates to be gradually integrated during inference (Gold and Shadlen, 2007; Hanks and Summerfield, 2017). Forming more precise estimates of the sensory world is often consequential for reinforcement, both in the lab (e.g. when a monkey categorises a stream of noisy sensory signals in return for liquid reward) and in the real world (e.g. when a consumer evaluates the quality of multiple relevant attributes of a product).

However, where information arrives in high volumes, the carrying capacity of the neural system can limit the precision of sensory estimates. Consider a binary discrimination judgment between two stimuli *A* and *B*, where information about their worth 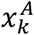 and 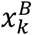 arrives in parallel via a sequence of *n* discrete sample pairs *k*. Let us assume that decisions are made by integrating and comparing transduced sensory estimates under the corrupting influence of “late” noise:

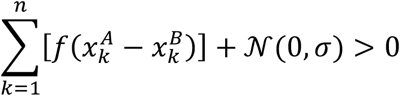

With limitless capacity (*σ* = 0) the best judgments will be made by simply comparing the summed estimates for A and B – in other words, the optimal *f*(·) is the identity function. Intuitively, if samples *x* are numbers and you are in possession of a calculator, the best thing to do is to simply compute their relative sum. However, under imperfect memory, comparative judgments of ground truth quality can paradoxically be promoted by more selective transduction that discards part of the sensory information (Tsetsos et al., 2016; Li et al., 2017; Spitzer et al., 2017; Moran and Tsetsos, 2018). For example, when *σ* > 0, a accuracy-maximising strategy is to selectively integrate the locally highest valued sample (i.e. 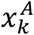 when 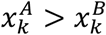) by partially down-weighting the less valued sample by a factor *w* (1 > *w* > 0). For illustration, imagine that deciding between two apartments to rent involves evaluating the candidates along multiple continuously-valued dimensions (price, location, size, etc). As memory demands become prohibitive (e.g. as *σ* grows), a strategy that is partly based on tallying the number of dimensions on which an alternative is higher valued (e.g. *w* < 1) will lead to more accurate choices. This is because ordinal (rank-based) strategies confer robustness on inference, just as nonparametric methods allow statistical calculations to be more robust to outlying data. Selective integration can be shown to be “optimal” because it optimises accurate responses given irreducible constraints on capacity. At the same time, however, selective integration is “irrational” in that it discards information and can lead to inconsistent choices, including violations of the normative axioms of transitivity and independence from irrelevant alternatives (Von Neumann and Morgenstern, 1944). Several models embodying this principle provide a good fit to human data in tasks involving sequential integration and comparison of discrete samples of information (Tsetsos et al., 2012, 2016; Bhatia, 2013; Summerfield and Tsetsos, 2015; Glickman et al., 2018; Gluth et al., 2018).

The framework of selective integration also makes predictions about neural signals, but these remain as yet untested. For example, several studies have shown that during sequential integration of decision information, the amplitude of single-trial EEG signals contralateral to the side of stimulation correlates with the strength of evidence conferred by each sample (Wyart et al., 2015). Thus, we might expect that those samples that are selectively integrated by virtue of being the local “winner” would be encoded with higher gain, i.e. a steeper linear relationship between *x* and relevant EEG signals. Here, we tested this view by recording scalp electroencephalographic (EEG) data whilst participants performed a task that involved averaging and comparing bar height within two parallel streams.

## MATERIALS AND METHODS

### Participants

Participants were 18 healthy adult volunteers with no history of neurological or psychiatric illness. Those who failed to complete both recording sessions (n = 2) or who performed at chance level (n = 1) were excluded. Analyses were conducted on the remaining 30 EEG sessions from 15 volunteers (female = 8; left-handed = 2; M_age_ = 24.29 ± 4.48). The study was approved by the Oxford University Medical Sciences Division Ethics Committee (MSD/IDREC/C1/2009/1) and informed consent was given at the start of each recording session. Monetary compensation was based on performance during the experiment (approximately £25 per participant).

### Experimental Design

All stimuli were created in the Psychophysics 3 Toolbox (RRID:SCR_002881; Brainard, 1997; Kleiner et al., 2007) for Matlab (MathWorks; RRID:SCR_001622). The experiment was presented on a 17-inch CRT monitor with 60 Hz presentation rate at a viewing distance of ∼70 cm.

On each trial participants viewed a sequence of 9 pairs of bars of variable height (Fig. 1A) that appeared in sequence, left and right of a central fixation dot. The task framing differed across sessions. In one recording session participants were asked to indicate with a button press the stream with the *highest* average bar height (‘high frame’) and in the other session they were asked to indicate the stream with the *lowest* average height (‘low frame’). The order of the two sessions was counterbalanced over participants and both sessions were separated by at least one week. We excluded participants who did not complete both sessions, because the framing manipulation allowed us to cleanly orthogonalize perceptual value (the raw bar height in pixels) from decision value (the level of evidence in favour of either response). The instructions stated that the bars indicated the value of two stock options whose prices fluctuated over time. They were asked to buy the most favourable (high frame) or sell the least favourable (low frame) stock option in different sessions.

**Figure 1.**
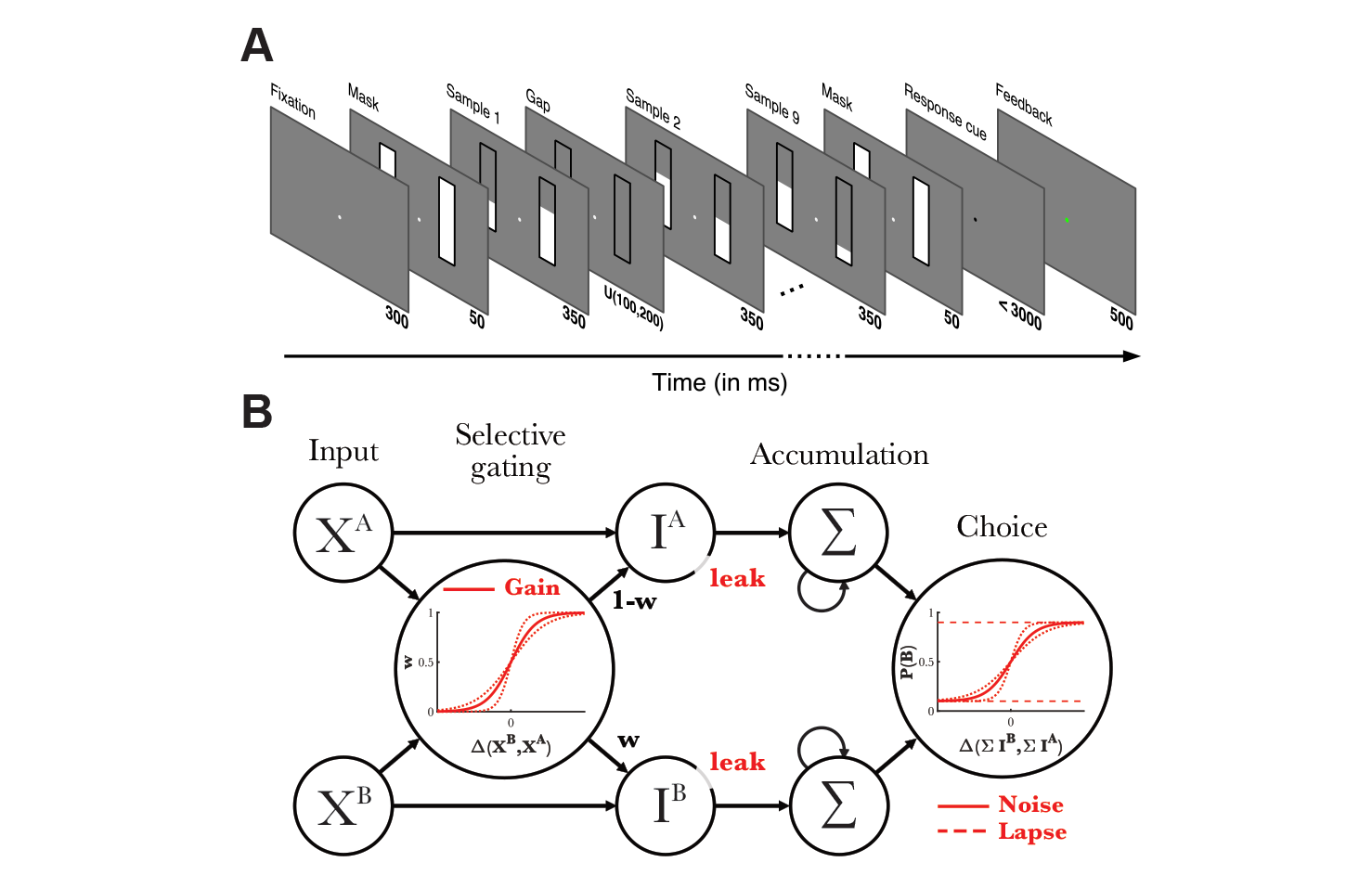
Task design and model schematic. **(A)** Participants viewed a sequence of nine pairs of bars varying in height and compared the average bar height of the left and right stream. In two separate recording sessions, participants indicated the sequence with either the highest or lowest average bar height. Numbers in bold below frames indicate duration of each frame in milliseconds. **(B)** The selective integration model assumes a separate accumulation process for each option, here the left and right bar sequence. Two concurrent input values *X*^*A*^ and *X*^*B*^ are transformed with a factor *w* based on their relative difference, down-weighting the lower input value and up-weighting the higher input value. All transduced values *I*^*A*^ and *I*^*B*^ are then integrated over time, with greater loss of information for earlier samples. Finally, the model generates a choice probability based on the difference in integrated values between the two streams.

Each trial started with a white fixation dot (radius = 5 pixels [px]) presented centrally on a grey background. The dot remained on screen for the whole duration of the trial. All bar stimuli were presented within two black rectangular placeholders (60 by 200 px; visual angle ∼4°) on either side of the fixation dot at a horizontal distance of 160 px. Each pair of bars remained on screen for 350 ms, and between successive pairs of bars, there was a jittered gap of empty black frames, lasting on average 150 ms (uniformly drawn between 100 – 200 ms). The first bar was a forward mask (300ms after fixation onset) of maximal bar height that occurred on both sides and was not included in the analysis. Bars were thus presented at a rate of ∼2 Hz (including the jitter) to minimise steady state neural responses. When all 9 bar pairs had been presented, a backward mask appeared again on screen for 50 ms. The fixation dot then turned black, indicating to the participants that they could respond by pressing the A and L keys on a QWERTY standard keyboard using the left and right hand to choose left and right stream respectively. Response mapping was fixed for the entire experiment. If participants failed to respond within 3 seconds, the fixation dot would turn red for 1000 ms and the words ‘Too late’ appeared in red above the fixation dot. When they responded before the deadline, the fixation dot would change colour for 500 ms: green for a correct response and red for incorrect. The next trial started after an inter-trial interval (ITI) of 300 ms. A trial could thus last for a maximum of 9350 ms (assuming 3 seconds with no response).

Trials were drawn from one of four bar height distributions (conditions), randomly intermixed throughout the experiment. In two conditions (‘Low variance’ and ‘High variance’), samples for the left and right stream were drawn pseudorandomly from a Gaussian distribution whose mean *μ* was in turn drawn from a uniform distribution between 80 px and 120 px, and whose standard deviation *σ* set to 10 (low variance) or 20 (high variance conditions). We used a resampling method to ensure that sample mean and variance closely matched the generative means and variance. To produce a correct answer, 6 px were subtracted from one of the two stream means.

We also added two more conditions which permitted greater control over the number of winning samples (i.e. those samples with locally greater decision evidence) on either side of the stream, allowing us to manipulate the correct answer independently from the number of winning samples contained within that stream. Samples for these two conditions, ‘Frequent winner’ and ‘Infrequent winner’, were generated as follows. In the frequent winner condition, the correct stream was manipulated to contain the winning sample in 2/3 of the sample pairs, while in the infrequent condition the correct stream only contained the winning sample 1/3 of the time. To achieve this, the mean bar height (*M*) of each trial was first drawn from a uniform distribution ranging between 110 and 130 px. Next, values were calculated separately for every three sample pairs (‘triplets’ *T*) in the trial. A deviation (*δ*) from the trial mean for each triplet pair *T* was drawn from a uniform distribution *∂* ∼ *U*(15,25). Additional noise *ε*_1_ and *ε*_2_ was drawn per triplet *T* from a normal distribution *ε* ∼ *N*(0,3). For every triplet *T*, the two streams *T*_*A*_ and *T*_*B*_ would have the following form:

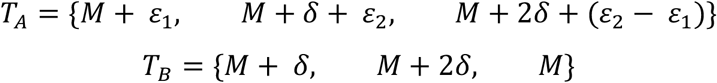

In this form, *T*_*B*_ always contains two winning samples (*M* + *δ* and *M* + 2*δ*), while the means of both triplets are identical. To produce a correct answer, 6 px were subtracted from one of the two stream means. In the frequent winner condition the mean of stream *A*, which contained most of the losing samples, was adjusted to make it the incorrect answer, while in the infrequent winner condition the mean of stream *B*, containing the most winning samples, was adjusted. Adjustments in low and high frames were inversions of one another.

To prevent participants from shifting gaze directly at one of the two streams, an area around the fixation dot (256 × 205 px; visual angle ∼2°) was defined in which participants had to keep fixation. Eye movements were monitored online using a Tobii EyeX eye-tracker (Tobii Technology, Stockholm, Sweden) and when the eyes moved outside the fixation area, the trial was classified as incorrect and the message ‘Eyes moved!’ appeared above the fixation dot. These trials were also omitted from analysis (0.35%).

Each recording session consisted of 600 trials, divided in 10 blocks, resulting in a maximum of 1200 trials (10 800 samples) for each participant. Participants could take self-timed breaks in between blocks. One session lasted around 2.5 h, including preparation of the cap, 60 minutes of task performance and removal of the cap at the end of the experiment.

### Selective integration model

The model decision is based on the output of two accumulators *Y*^*A*^ and *Y*^*B*^ that integrate the input values of the left and right stream respectively (Fig. 1B). Input values *X*^*A*^ and *X*^*B*^ are the raw pixel heights *H* of each sample (bar), inverted for the low frame (200 – *H*). Each integrator is updated separately for each sample *k* according to the following formula:

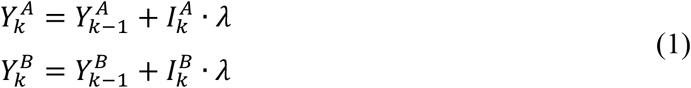

*Y*^*A*^ and *Y*^*B*^ are both initialized to zero. *I*^*A*^ and *I*^*B*^ are the transformed values of the input after graded selective integration:

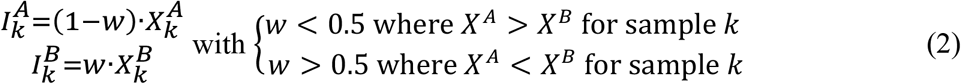

Where *w* is a gating variable that is determined by passing the difference between input values (Δ*X* = *X*^*B*^ − *X*^*A*^) through a logistic function

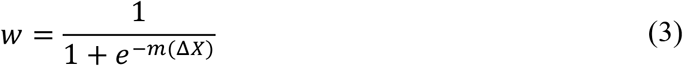

with the slope of *m* determining the extent to which the difference between the two input values attenuates the input values *X*. Values of *m* close to zero indicate a limited sensitivity to the difference in bar height, treating the gating of locally ‘winning’ and ‘losing’ samples almost equally when the difference between them is small versus when their difference is large (note that the ‘winning’ sample changes with the frame of reference i.e. whichever sample is highest in the high frame and lowest in the low frame). Larger values of *m* then indicate an increased sensitivity to the magnitude of the difference between the bar heights in a sample pair, giving rise to a tendency to more strongly overweight the locally winning sample for larger differences; negative values of *m* have the converse effect. The transfer function itself ensures that for very large differences between the bars, where it is clear which of the two bars wins, the winning input value is integrated close to its original value, while the losing sample value is almost entirely suppressed (i.e. w is close to 0 or 1). On the other hand, for input values that lie close together and comparison is difficult, both values carry approximately equal weight to their accumulators. A recent study demonstrated that a ‘graded’ selective integration such as the one proposed here outperformed a ‘binary’ selective integration that works independent of the size of the sample difference, both in predicting human choices and its robustness to noise (Glickman et al., 2018).

The model further assumes a leaky accumulation, with earlier samples carrying less weight on the final decision. Each transduced sample is therefore multiplied with the leak function value:

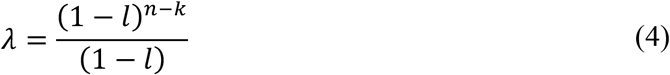

with *l* the amount of information loss and *n* the total number of samples.

Finally, the model outputs a choice probability based on the difference between the accumulators after all evidence has been accumulated:

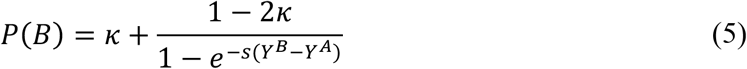

*s* is the slope of the response function, often referred to as ‘late’ or ‘integration noise’, a factor that adds uncertainty to the final choice. Finally, a lapse rate parameter *κ* was added to the response function to allow for higher error rates in the model.

### Model fitting procedure

The model has four free parameters: selective gating parameter *m*, leak *l*, late noise *s* and lapse rate *κ*. Best fitting parameter sets for each participant were obtained by minimizing the negative sum of the log-likelihoods using a scatter-search based global optimization solver in Matlab. The parameter search space was constrained as follows:

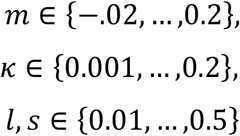

To test whether the gating parameter *m* significantly contributed to the model performance, two models were compared: the full SI model and a fixed gating model. For the fixed gating model *m* was set to 0, which effectively gave equal weight to each input (*w* = 0.5), whereas for the full model *m* was unrestricted, allowing *w* to take any value between 0 and 1. Model fits were compared through split-half cross-validation, a method that allows comparing models with different numbers of free parameters through their generalizability to new data. One recording session would serve as the training set and the best fitting parameters obtained from the training set were then used to estimate how well they predicted responses in the other session and vice versa. Log-likelihoods from both test sets were then summed per participant and Wilks’ test was used to identify the most appropriate model. Best fitting parameters for the model used in subsequent analyses were estimated from the collapsed data over both recording sessions.

### Parameter recovery

To ensure there was no trade-off between the parameters of our model, we tested whether simulated parameter combinations could be recovered using the same model-fitting procedure as above. We were mainly interested in the possible trade-off between the *m* and *s* parameters and therefore performed our parameter recovery procedure once for each parameter of interest. First, we generated 25 equidistant values over the full sampling range of the parameter of interest. We then generated 500 random combinations of the three remaining parameters. The sampling space of all parameters was limited by the minimum and maximum estimated values from the real data. At every step of the parameter of interest, model responses were generated for the 500 parameter combinations using the input data from a randomly selected subject each time (1200 trials). Finally, the input data and model generated responses were used to estimate best-fitting parameters again and test how well we could recover the original parameter combinations. Statistical tests on the recovery error for the remaining three parameters were performed on the 25 values obtained after averaging over the 500 random iterations at each step.

### Statistical analysis

All analyses of human data were performed on the collapsed data from both recording sessions, with input values for the ‘low frame’ session inverted (200 − *H*) to reflect the momentary decision value *X* rather than raw bar height *H*. The impact of winning and losing samples (i.e. the bars with respectively the highest and lowest decision evidence in a sample pair) on choice was tested through a generalized linear model (GLM), modelled with a binomial distribution and a logit link function, predicting the probability of choosing the right stream, as follows:

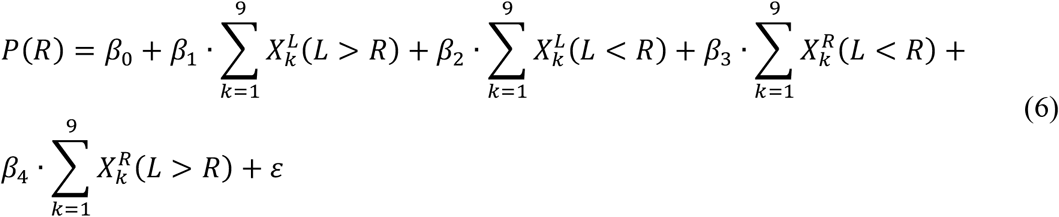

The four parametric regressors in the model coded for the cumulative sum of input values for all (1) winning samples on the left side [*X*^*L*^(*L* > *R*)], (2) losing samples on the left side [*X*^*L*^(*L* < *R*)], (3) winning samples on the right side [*X*^*R*^(*L* < *R*)] and (4) losing samples on the right side [*X*^*R*^(*L* > *R*)]. The regressors were always standardized before parameter estimation. Parameter estimates for the left stream were then sign-flipped and averaged with the estimates of the right stream, in order to obtain an average modulation of winning and losing samples.

The SI model further allows for a ‘recency effect’ through the leak parameter, whereby earlier samples carry less weight in the final choice than later samples. We tested this assumption through a new GLM predicting choices for the right stream based on the signed difference of each sample pair (right minus left).

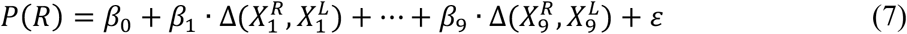

The model thus consisted of 9 parametric regressors representing the serial position in the stream. If choices were driven more by more recent samples, coefficient estimates should be higher for regressors coding for later samples.

All statistical tests were performed at the group level. Given the relatively low number of subjects, we opted for non-parametric tests that do not assume a normal distribution of the data.

### EEG acquisition and pre-processing

EEG signals were recorded using a Neuroscan system with SynAmps-2 digital amplifiers with 60 Ag/AgCl electrodes located at FP1, FPz, FP2, F7, F5, F3, F1, Fz, F2, F4, F6, F8, FT7, FC5, FC3, FC1, FCz, FC2, FC4, FC6, FT8, T7, C5, C3, Cz, C2, C4, C6, T8, TP7, CP5, CP3, CP1, CPz, CP2, CP4, CP6, TP8, P7, P5, P3, P1, Pz, P2, P4, P6, P8, PO7, PO5, PO3, POz, PO4, PO6, PO8, O1, Oz, and O2. Four additional EOG electrodes in dipolar montage (two horizontal, two vertical) were recorded, together with one mastoid for reference. Electrode impedance was brought below 10 kΩ before recording. Data was collected at a sampling rate of 1 kHz and high-pass filtered at .01 Hz.

Pre-processing was done in Matlab using functions from the EEGLAB toolbox (RRID:SCR_007292; Delorme and Makeig, 2004) and custom scripts. First, data was downsampled to 250 Hz, subsequently low-pass filtered at 40 Hz and then high-pass filtered at 0.5 Hz. Excessively noisy channels were identified through visual inspection for each participant and interpolated based on the weighted average of the surrounding electrodes. Next the data was re-referenced offline to average reference (excluding EOG channels). Trial epochs were extracted spanning 1 second prior to the fixation dot onset until 7 seconds after. Epochs were subsequently baselined relative to the pre-fixation time window of −500 to 0 ms. Artefacts related to eye-blinks and other sources of consistent noise were identified through Independent Component Analysis (ICA) and removed from the data after visual inspection. Finally, the data were epoched again at different times for various analyses. For the sample-based regressions, the data were epoched relative to each sample pair onset, starting at 100 ms before until 750 ms after sample onset and baselined again relative to the full pre-stimulus window, to exclude any systematic offset during the length of the trial. For response-locked analyses, the epoch was set to 3 seconds prior to response onset to 300 ms after. The baseline window was chosen at 3 to 2.5 seconds before response onset.

HEOG channel data were pre-processed using a similar pipeline, apart from average referencing and ICA.

### Time-frequency transformation

The pre-processed epochs spanning 8 seconds were transformed into the time-frequency domain using functions from the Fieldtrip toolbox (RRID:SCR_004849; Oostenveld et al., 2011) for Matlab and custom scripts. Power was calculated every 25 ms within an epoch by convolving a sliding Morlet wavelet (width = 7 cycles), for frequencies between 8 and 38 Hz, incremented in steps of 3 Hz. To allow for a comparison of power estimates over frequency bands, each frequency within an epoch was dB transformed relative to the baseline window [−300 300] ms, a ‘silent’ period including the ITI and start of a new trial, using the following formula:

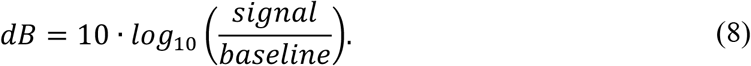

The data was subsequently epoched into smaller epochs between [−100, 750] ms relative to sample pair onset. For visualization purposes the time-frequency results were interpolated, while cluster statistics were performed on the raw data.

### EOG control analysis: regression model

Although we tried to control for large saccades with the eye-tracker, there remained the possibility that small eye movements towards the winning sample were driving some of our neural findings. Because we did not have access to very precise eye-tracking data as it was only used for online monitoring, we turned to the signals recorded with the horizontal dipolar electrodes (HEOG). To assess how eye movements were influenced by decision information, here operationalised as the difference in momentary evidence, we constructed a regression model with the epoched HEOG signal as the dependent variable.

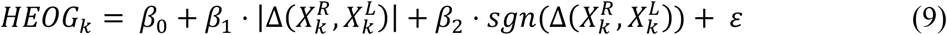

The difference in decision information 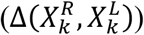 was split into two regressors: the absolute strength of evidence and the sign (or direction) of the information. The regression was then performed for each sample epoch independently at each time point and electrode, identical to the single-trial EEG regression.

### EEG analysis: regression models

To understand how EEG signals were modulated by the stimulus information, we used a regression-based approach. Single-trial EEG data was regressed against parametric predictors for each time-point and electrode independently. This analysis tests for a linear relationship between the magnitude of decision values and the momentary amplitude of the EEG signal at each timepoint following each sample. Coefficient estimates obtained from the regression model reflect the slope of the modulation, i.e. the strength of the linear relationship. Group-level statistics were subsequently computed over the obtained time courses of parameter estimates over all participants.

First, we constructed a sample-based linear regression model that allowed us to dissociate the neural modulation of winning and losing samples:

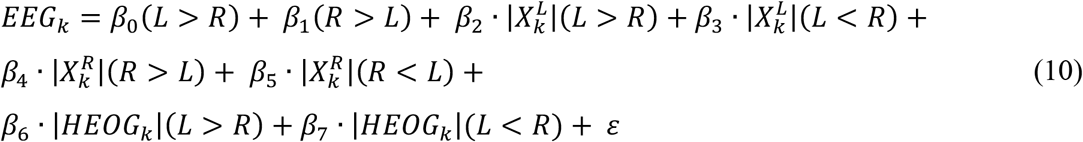

with *β*_0_ and *β*_1_ as binary indicator variables for samples where the left (L) or right (R) bar in the pair had the highest decision value respectively. The four parametric regressors *β*_3_ - *β*_6_ code for the absolute decision evidence (*X*_*k*_) of the bar when the left bar won (*β*_3_), left lost (*β*_4_), right won (*β*_5_) and right lost (*β*_6_). Effectively, each sample pair related to four regressors (*β*_0_, *β*_1_ and two of the four parametric regressors). The trial mean was subtracted from all bars to eliminate any differences related to the absolute size of the bars. The HEOG signal was added as a nuisance regressor to account for variability related to small eye movements. The single trial values of the HEOG channels were calculated based on an averaged time window of 100 ms centred around the negative peak of the parametric regressors (176 - 276 ms) from an initial model that did not contain the HEOG nuisance regressors. Single-trial EEG signal locked to stimulus onset was subsequently regressed against the full model with 8 regressors, independently for each sample pair. Estimated time courses were averaged over all sample pairs before statistical inference.

We defined two regions of interest (ROI) based on previous research (Wyart et al., 2015) to test for early modulation of sample evidence: left (O1, PO5, PO7, P5, P7, CP5, TP7) and right (O2, PO6, PO8, P6, P8, CP6, TP8) occipito-parietal (OP) electrodes. The coefficients *β*_0_ and *β*_1_ in our regression model encode the average EEG signal for sample pairs where the left (*β*_0_) or right sample won (*β*_1_). To compare categorical modulation of stimulus-evoked activity, we averaged the activity over those electrodes contralateral to the winning sample, namely right OP electrodes for *β*_0_ (trials where the left sample won) and left OP electrodes for *β*_1_(trials where the right sample won). This is equivalent to flipping the scalp maps and treating each sample pair as if the right sample won. The opposite hemispheres then contained the neural signals modulated by the losing sample. Time series for a parametric modulation of winning samples (X_win_) were obtained in a similar way: averaging over the right OP electrodes when left won (*β*_3_) and left OP electrodes when right won (*β*_5_). Time series for parametric modulation of losing samples (X_lose_) were obtained at right OP electrodes when left lost (*β*_4_) and left OP electrodes when right lost (*β*_6_). Significance of adjacent time points in the averaged time series were tested using nonparametric cluster-based permutation tests (cluster-defining threshold and corrected significance level at p < 0.05, 5000 iterations) (Maris and Oostenveld, 2007). Estimated time series were smoothed per subject using a low-pass filter (Butterworth filter, cut-off = 25 Hz), removing small fluctuations in the signal that could obscure larger clusters from the permutation test.

The same regression model and contrasts were used to study differences in lateralization of alpha band activity, with an additional iteration over frequency bands. The averaged time window was temporally smoothed with a 100 ms full-width at half-maximum (FWHM) Gaussian kernel before cluster-based permutation test (cluster-defining threshold and corrected significance level at p < 0.025). We used a lower p-value threshold for our cluster correction algorithm for the time-frequency analysis, to avoid clusters from different frequency bands to merge into one, potentially inflating the reliability of smaller clusters, a well-known problem with cluster-based analysis methods (Maris & Oostenveld, 2007).

Next, we sought to identify neural signals related to evidence accumulation. We formalized ‘accumulated evidence’ as the difference between the cumulative sum of evidence of the right minus left stream. The model tested evidence integration during the course of the trial and was therefore conducted at the sample level (for samples 2 to 9), regressing EEG signals from sample *k* against the absolute sum of differences up to the previous sample (i.e. 1 to *k* − 1) and controlling for the evidence on sample *k*:

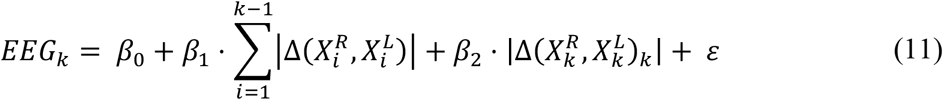

Since the outcome is rectified, i.e. unsigned with respect to the response hand, this measure was not confounded with motor responses. Because the cumulative difference for early samples was too highly correlated with the generative data, making it hard to disentangle the two, we opted to derive the independent variables from the model estimates rather than the generative data.

## RESULTS

### Behavioural results

We first examined the behavioural data to test whether participants weighted the samples as described by the selective integration model. We fitted a logistic regression model that separately estimated the influence of winning and losing samples on right-hand choices (Eq. 6). The null hypothesis is that there will be no difference in how much influence winning samples (e.g. *X*^*A*^ when *X*^*A*^ > *X*^*B*^) and losing samples (e.g. *X*^*B*^ when *X*^*A*^ > *X*^*B*^) carry on choice. However, we found that parameter estimates for winning samples (Mdn *β*_*win*_= 3.12) were significantly higher than for losing samples (Mdn *β*_*lose*_ = 3.06; Z = 3.41, p < 0.001, two-sided Wilcoxon paired signed rank test), suggesting there is indeed a difference in how samples were weighted according to their relative decision value (Fig. 2A).

**Figure 2.**
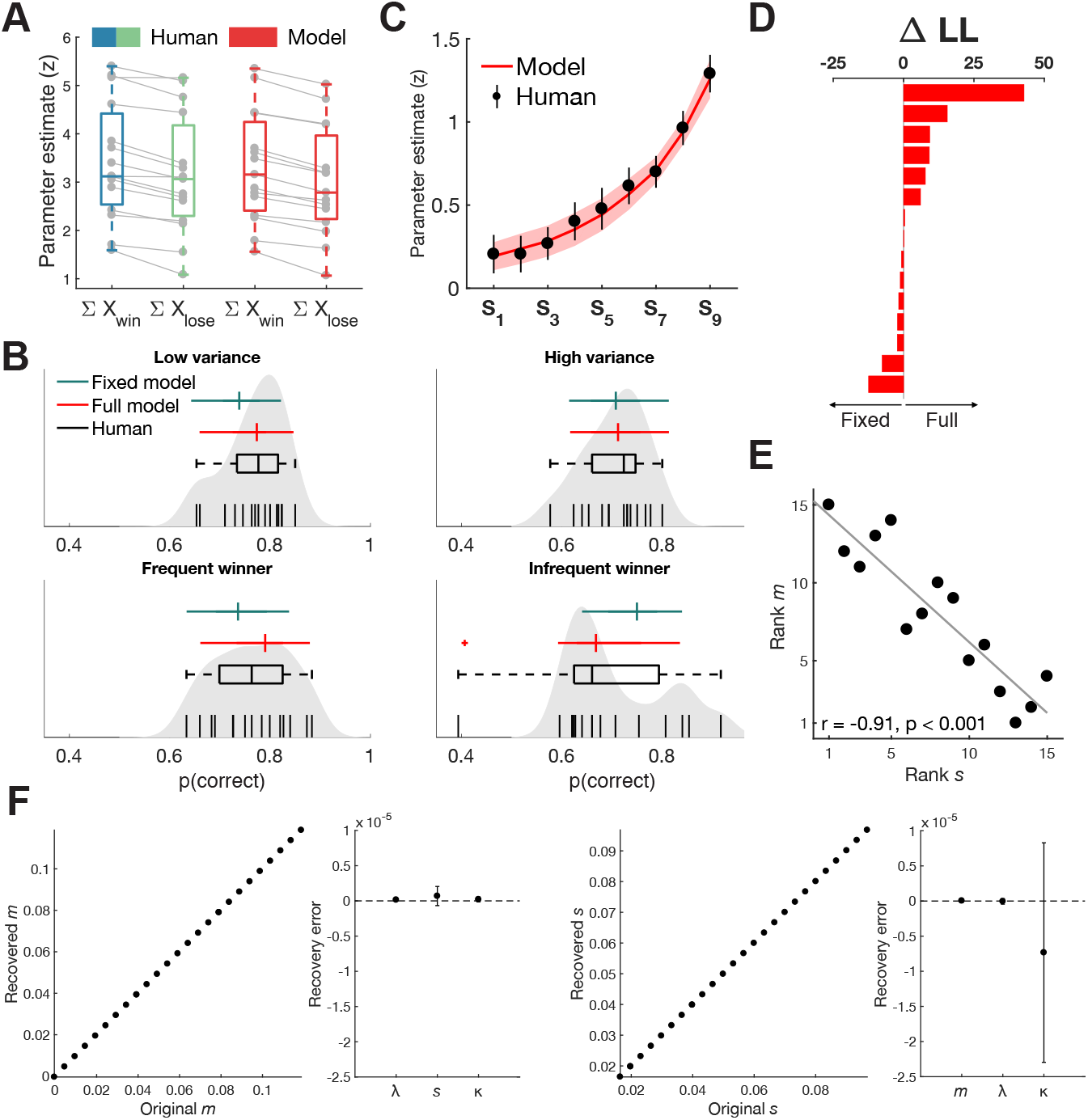
Behavioural results and model fits. **(A)** Parameter estimates from a regression model predicting the probability of choosing the right-hand sequence, both in humans (blue/green) and the fitted SI model (red). In line with predictions of the SI model, sample values were weighted higher on average when decision evidence was greater relative to the concurrent sample. A SI model fitted to the choice data was able to recreate this pattern. **(B)** SI predicts that a tendency of tallying winning samples will lead to lower accuracy in trials where the incorrect sequence contains most winners. Performance was indeed lower in the ‘infrequent winner’ condition, but not significantly so from the ‘frequent winner’ condition. Qualitatively, the full SI model (red) resembled human accuracy better in all four conditions compared to a model where the gating parameter was fixed to 0. Black vertical lines show individual participant’s accuracy and grey shaded area represents non-parametric kernel estimate (bandwidth = 0.03) of the accuracy distribution in our sample. **(C)** Parameter estimates from a new regression model showed a ‘recency effect’, where earlier samples carried less weight on participants’ final choice. The fitted SI model closely followed the same trend (red). Error bars and shaded area represent 95% confidence interval. **(D)** Individual differences in LL estimates (full – fixed) after cross-validation. **(E)** The slope of the response function (*s*) in the fitted SI model correlated significantly with the slope of the selective gating function (*m*), suggesting participants with greater levels of late noise compensate through stronger gating of information. Grey line represents least-squares fit line. **(F)** Parameter recovery error was not significantly different from zero, both when systematically varying *m* (left two panels) or *s* (right two panels). Error bars represent 95% confidence interval.

If participants give more weight to winning samples, their performance might be poorer in a condition where the incorrect stream contains the most winning samples, a so-called ‘frequent-winner’ effect (Tsetsos et al., 2016). To maximize the opportunity to compare trials where the winning stream contained the most versus the fewest winning samples, we included two conditions that explicitly contained these types of trials (‘frequent winner’ and ‘infrequent winner’) as well as two conditions where the height variance was either low or high. A non-parametric omnibus test (Friedman test) first confirmed there were differences in choice accuracy between the four conditions (*X*^2^ = 13.34, p = 0.004; d.f. = 3, N = 15) (Fig. 2B). We then ran a post hoc pairwise comparison between all conditions (two-sided Wilcoxon signed rank test) and corrected for multiple comparison using FDR correction (Benjamini and Hochberg, 2009). Unlike previous reports (Tsetsos et al., 2016), we did not observe a statistically significant difference between the ‘infrequent winner’ (Mdn = 0.66) and ‘frequent winner’ conditions (Mdn = 0.76; Z = 1.62, p = 0.1054), although the results did show a numerical trend in the predicted direction and a selective integration model fitted to the human choice data predicted a difference between the (in)frequent winner conditions (see below). Participants performed significantly worse in the ‘infrequent winner’ condition, but only in comparison to the ‘low variance’ condition (Mdn = 0.77; Z = 2.24, p = 0.0248). Participants also performed significantly worse in the more difficult high variance condition (Mdn = 0.72) compared to the low variance condition (Z = 3.14, p < 0.001).

Finally, the SI model predicts a ‘recency effect’, where samples presented later in a trial should carry more weight on the final decision, because there was less time for this information to be lost. A new logistic regression model predicting right-hand choices (Eq. 7) showed that indeed, when sample pairs were assessed based on their serial position in a trial, parameter estimates increased over time, indicated by a significantly increasing slope fitted to each subject’s parameter estimates (Fig. 2C; Mdn slope = 0.13; Z = 3.41, p < 0.001).

### Model fits to human data

A more formal test for selective integration can be achieved by fitting the SI model to human data and comparing it to an equivalent alternative model that does not show selective integration. A full SI model with 4 free parameters (gating *m*, leak *l*, late noise *s* and lapse rate *κ*) was fitted to each participant’s trialwise choice data (*m* = 0.0377 ± 0.03; *l* = 0.234 ± 0.1; *s* = 0.0414 ± 0.03; *κ* = 0.0176 ± 0.02; Table 1). To assess whether the gating parameter meaningfully contributed to the model fits, we compared the full model to a fixed gating model where the gating parameter *m* was fixed to 0, eliminating the unequal weighting of sample inputs. Both models were tested through two-fold cross-validation, estimating parameter fits on one recording session and testing model predictions on the other session and negative log-likelihood (LL) estimates of the test sets were summed per participant. The full model (mean LL = −623.57) was favoured over the fixed model (mean LL = −627.77; *D* = 8.39, p = 0.0038, Wilks’ test) suggesting the gating parameter of the SI model meaningfully contributed to explain choice behaviour (Fig. 2D). To determine qualitative fits of both models, we reran our behavioural analyses on model choices. The full model was able to reproduce the difference in parameter estimates for winning and losing samples (Fig. 2A) and the recency effect (Fig. 2C), even though the model was not specifically fit to these data points. Only the full model was further able to capture the lower performance in the ‘infrequent winner’ condition (Fig. 2B), and in general captured the patterns in human performance per condition qualitatively better than the fixed model.

**Table 1.** Model fitting results. Individual parameter estimates from the full model and negative log-likelihood estimates for the cross-validated fixed and full model.

| Participant | $m$ | $\lambda$ | $\kappa$ | $s$ | LL<sub>fixed</sub> | LL<sub>full</sub> |
| --- | --- | --- | --- | --- | --- | --- |
| 1 | 0.0004 | 0.1191 | 0.0000 | 0.0935 | -526.54 | -527.72 |
| 2 | 0.0572 | 0.2119 | 0.0386 | 0.0205 | -658.74 | -643.22 |
| 3 | 0.0509 | 0.1684 | 0.0675 | 0.0186 | -698.88 | -711.32 |
| 4 | -0.0004 | 0.1384 | 0.0000 | 0.0909 | -598.25 | -599 |
| 5 | 0.0333 | 0.2902 | 0.0000 | 0.0343 | -567.93 | -575.55 |
| 6 | 0.0035 | 0.3063 | 0.0603 | 0.0968 | -670.1 | -671.84 |
| 7 | 0.1187 | 0.3260 | 0.0000 | 0.0164 | -707.01 | -664.21 |
| 8 | 0.0544 | 0.3134 | 0.0120 | 0.0198 | -669.34 | -661.61 |
| 9 | 0.0397 | 0.4236 | 0.0000 | 0.0242 | -679.32 | -673.31 |
| 10 | 0.0014 | 0.1171 | 0.0267 | 0.0756 | -609.2 | -611.42 |
| 11 | 0.0395 | 0.0827 | 0.0260 | 0.0235 | -535.95 | -526.81 |
| 12 | 0.0501 | 0.2124 | 0.0227 | 0.0286 | -571.56 | -562.29 |
| 13 | 0.0532 | 0.1652 | 0.0000 | 0.0181 | -628.48 | -628.32 |
| 14 | 0.0202 | 0.3191 | 0.0000 | 0.0319 | -687.96 | -687.57 |
| 15 | 0.0429 | 0.3155 | 0.0096 | 0.0316 | -607.33 | -609.48 |

Finally, Tsetsos et al. (2016) showed that the fitted parameters for gating and late noise were highly correlated in their data set (but not for simulated data), suggesting that the strength of the gating process compensates for higher levels of integration noise. We replicated this finding in our data set (Spearman *ρ* = −0.91, p < 0.0001) (Fig. 2E). To ensure that this correlation was not due to a trade-off between the two parameters (*m* and *s*), we performed parameter recovery on simulated parameter combinations while systematically varying either *m* or *s* and assessed how well our fitting procedure could recover the original parameters. As shown in Figure 2F, we could successfully recover the original parameters, while the recovery error was not significant for any of the remaining parameters (all p > 0.115). Moreover, we could not find a correlation between *m* and *s* in either recovery procedure (all r < 0.0001, all p > 0.99). The larger confidence interval for *κ* when varying *s* possibly indicates a trade-off between the slope and lapse rate of the response function.

### Eye movement control analysis

Despite our best attempts to control for large eye movements, there remains the possibility that smaller eye-movements occurred undetected and influenced our neural findings. To better understand how decision information could be driving eye movements, we decided to run a control analysis directly explaining trial-by-trial fluctuations in horizontal eye movements. We regressed dipolar HEOG signals against the absolute difference in decision evidence and the sign (i.e. direction) of the difference (Eq. 9). We found that indeed eye movements were significantly explained by the sign of the decision information between ∼300 and 500 ms (p_cluster_ < 0.05; Fig. 3), but not by the strength of the evidence. In other words, the eyes would move towards the winning sample. Importantly, as we will see below, these effects only arose around 300 ms and peaked at ∼400 ms, after the observed parametric modulation of decision evidence in contralateral electrodes. This suggests that the modulation of posterior EEG signals is unlikely to be driven by small shifts in eye movements. Nevertheless, we decided to include HEOG channel data as a nuisance regressor in subsequent analyses to exclude the possibility of eye movement contamination.

**Figure 3.**
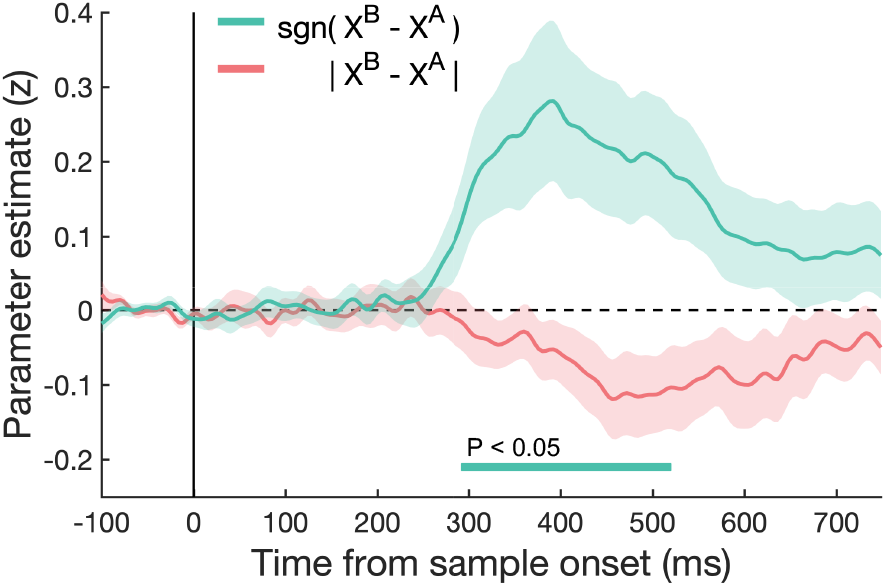
Eye movement control analysis. When regressing HEOG channel data against the momentary evidence, the sign but not the strength of decision evidence, significantly predicted eye movements starting around 300 ms. Crucially, these effects arose later than our lateralized neural findings. Shaded coloured area indicates SEM.

### Early modulation of posterior EEG activity

The SI model states that selective gating occurs at the level of the individual samples before evidence is passed to a subsequent integration stage. We thus predicted that posterior EEG signals would encode locally “winning” samples with higher gain than “losing” samples, i.e. that the slope of regression linking decision information to EEG amplitudes would be steeper for winners than losers. In addition to this multiplicative effect, we also tested for an additive bias, i.e. that EEG signals were higher contralateral to the winning sample, irrespective of its input value.

Since samples were presented parafoveally, we expected these effects to occur contralateral to the location of each sample. We therefore focused our analyses on two a priori defined posterior regions of interest (ROI): left and right occipito-parietal electrodes. We constructed a linear regression model (Eq. 10) with two intercepts, coding whether the left or right sample won within a sample pair (to test the additive effect), and four parametric regressors coding the sample evidence separately for left and right and whether the sample won or not (to test the multiplicative effect).

We first examined the coefficients associated with the parametric regressors that, at each time point and electrode, reflect the slope of the relationship between the magnitude of the decision evidence and EEG amplitude, separated for winning and losing samples. After collapsing over posterior electrodes contralateral to winning or losing samples, we found that the sample evidence of both winning and losing samples was encoded in the EEG signal around 250 ms, but more importantly, this was significantly stronger for winning samples (p_cluster_ < 0.05; Fig. 4A). This suggests that, as predicted, samples carrying equal decision evidence are encoded more strongly in the EEG signal when they are the winning sample in a sample pair.

**Figure 4.**
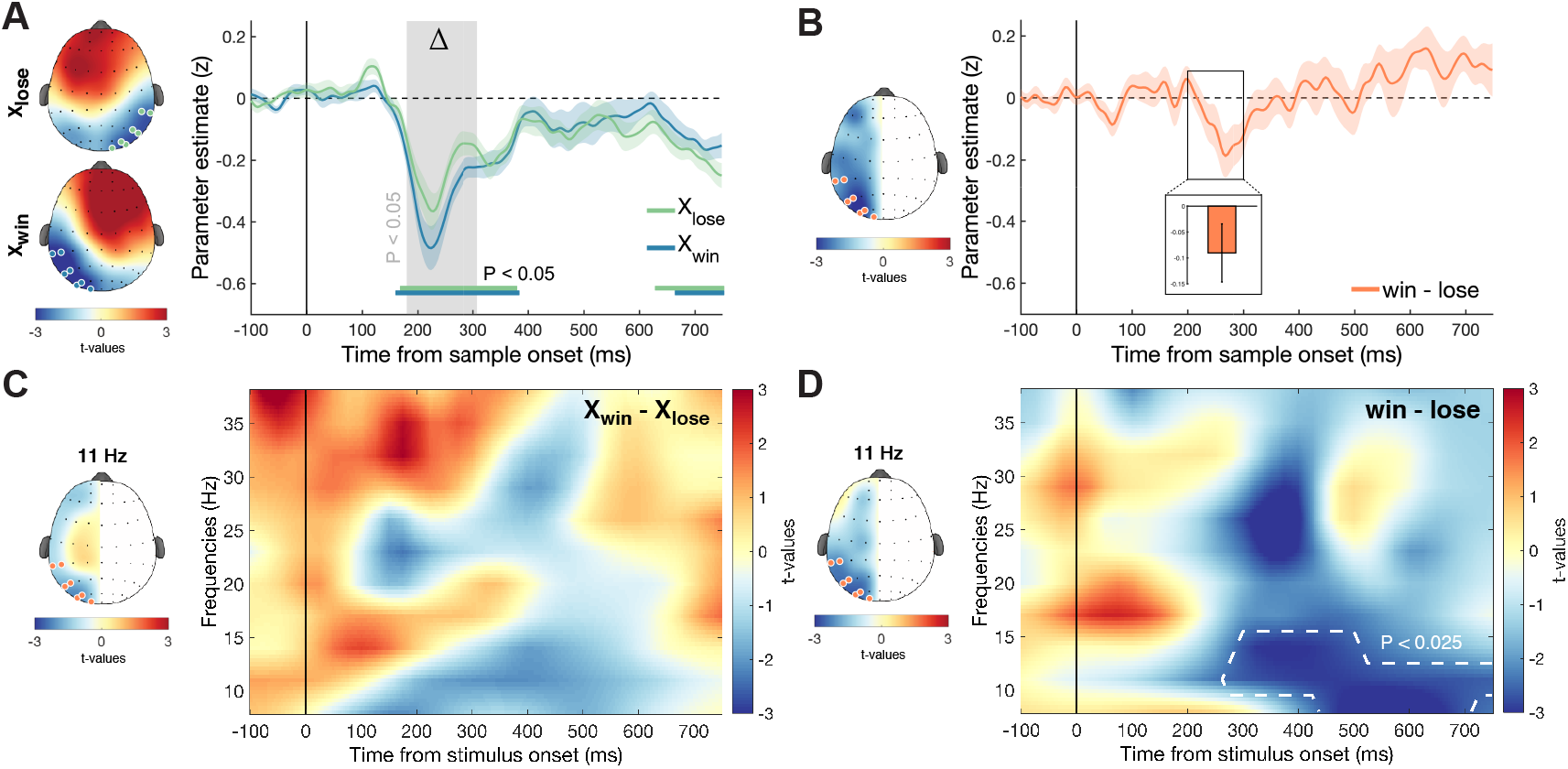
Early modulation by relative sample evidence in posterior electrodes. A linear regression model was used to separately assess the influence of winning and losing samples on EEG signals at each time point, electrode and sample pair. Coefficients for left sample regressors in our model were horizontally flipped, so winning samples were projected to the left hemisphere and losing samples to the right hemisphere. Plots represent coefficient estimates averaged for all nine sample pairs. **(A)** Coefficient estimates for the parametric regressors in our model showed that winning and losing samples were both significantly encoded in the EEG signal around 250 ms (bottom coloured lines, p_cluster_ < 0.05) in posterior electrodes (coloured dots on the scalp plots) contralateral to the sample location. More importantly, winning samples were encoded significantly stronger compared to losing samples (grey shaded area, p_cluster_ < 0.05). Shaded coloured area indicates SEM. Top and bottom scalp plot show significance of coefficient estimates for losing and winning samples respectively at the identified time window of significant dispersion. **(B)** By contrasting the intercept coefficients of the regression model, we observed a small additive effect that was stronger for winning samples compared to losing samples, but only when averaging between 200 and 300 ms after stimulus onset in posterior electrodes. The orange line represents the difference between the time series of the two intercepts. Scalp plot shows the left minus right hemispheric difference of coefficient estimates. Shaded coloured area represents SEM. **(C)** An identical regression model was used to explain time-frequency data. We did not find a significant difference in coefficient estimates between winning and losing samples based on their parametric values. Colour map shows difference between winning and losing samples at the electrodes indicated on the scalp plot. Scalp plot shows difference between winning and losing samples and hemispheres at same time window as (D) for the 11 Hz frequency band. Time-frequency data was interpolated for visualization purposes, while cluster statistics were performed on the raw data. **(D)** However, we did observe an additive effect in alpha frequencies (8 – 12 Hz), with greater suppression for winning samples compared to losing samples in the same contralateral posterior electrodes starting around 375 ms after sample onset (p_cluster_ < 0.025). Colour map shows the difference between winning and losing samples at the electrodes indicated on the scalp plot. Scalp plot shows difference between winning and losing samples and hemispheres at the time of the identified cluster for the 11 Hz frequency band.

The intercepts of the regression model then reflect the average signal in trials where either the left or the right sample won, independent of their decision evidence. At each time point and electrode, the parameter estimates reflect an additive effect of sample identity (winner or loser) on EEG signals. To test a difference in the additive effect of winning and losing samples, we averaged intercept parameter estimates at ROI electrodes contralateral to all winning samples and compared them to the averaged intercept parameter estimates at ROI electrodes contralateral to all losing samples. This initially yielded a significant effect whereby signal in posterior electrodes responded to winning samples more strongly than losing samples (p_cluster_ < 0.05; Fig. 4B) between ∼200 and 300 ms after sample onset. However, when adding HEOG nuisance regressors, the effect was attenuated and no longer survived cluster correction (p > 0.05). It did, however, remain robustly significant when averaging over a 200 – 300 ms time window (Z = −3.85, p = 0.0001, Wilcoxon signed rank test). We thus interpret the additive effect with caution; there is a possibility that small eye movements could at least partially explain the neural effects we find here.

The negative contralateral modulations reported here show a strong resemblance to the well-known N2pc component from event-related potential (ERP) literature (Luck and Hillyard, 1994; Luck, 2012), a posterior negativity contralateral to the (covertly) attended target starting around 200 ms after stimulus onset. For the additive effect, this is expected, because the analysis we conducted on the intercept is closely related to the computation of an ERP. However, the fact that this signal is also modulated by the decision information nuances and extends previous work with trial averages.

### Alpha suppression

It has previously been suggested that SI could occur when selective attention is oriented to the winning sample (Glickman et al., 2018). Previous work has reported that a lateralized suppression in alpha (8 – 12Hz) power occurs when participants orient visuospatial attention towards a contralateral target (Sauseng et al., 2005; Bacigalupo and Luck, 2019). We used the same sample-based regression model as above (Eq. 10), but now predicting sample-wise fluctuations in power over frequency bands between 8 and 38 Hz. The same occipito-parietal contrasts for the intercepts of the model revealed a significant cluster of greater alpha suppression contralateral to the winning sample compared to the losing sample from ∼375 ms onwards (p_cluster_ < 0.025) (Fig. 4D). Note that these effects survived the inclusion of horizontal eye movements and might imply that covert attention was dynamically shifting to the winning sample. We could not find any statistical evidence for a parametric modulation of alpha suppression by input value (p_cluster_ > 0.05) (Fig. 4C).

### Evidence accumulation

In the SI model, samples are first weighted according to their relative value, and then integrated into a cumulative signal that eventually determines choice. Previous studies have observed in both perceptual and value-based decision making tasks that central EEG signals build up in concert with the strength of accumulated evidence (Donner et al., 2009; O’Connell et al., 2012; van Vugt et al., 2012; Twomey et al., 2015; Pisauro et al., 2017; von Lautz et al., 2019). We examined encoding of accumulated evidence over the course of the trial with a regression model that tested whether the cumulative difference in absolute (unsigned) decision information up to sample *k* − 1 was encoded in neural signals following the onset of sample *k* (Eq. 11). Accumulated evidence was significantly encoded around 250 – 400 ms in the proximity of central electrode Cz (p_cluster_ < 0.05; Fig. 5) suggesting neural signals were tracking the strength of evidence up to the current sample. Although the focus of these signals is slightly more frontal than previously reported, they resemble the “centro-parietal positivity” or CPP reported by O’Connell et al. (2012), which has also been related to the updating of the decision evidence.

**Figure 5.**
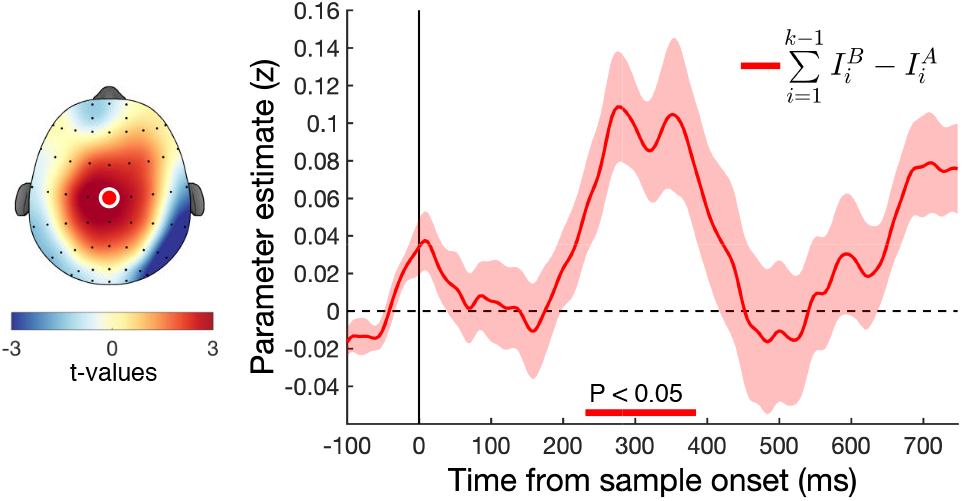
Encoding of evidence integration in central electrodes. We found a significant positive relationship with the accumulated evidence of 1: *k* − 1 in central electrode Cz around 300 ms after stimulus onset. Scalp plot shows EEG activity averaged over the significant cluster around 300 ms. Bottom coloured lines indicate significant clusters (p_cluster_ < 0.05). Coloured shaded area indicates SEM.

## DISCUSSION

In this report we tested for, and obtained, neural evidence in support of a “selective integration” policy during human decision-making. Input values conferred by each of two samples (bars) was weighted differently according to its local rank, and this behavioural effect was accompanied by modulation of neural signals over posterior electrodes. This modulation arose early after sample onset (∼ 250 ms), while signals encoding the strength of accumulated evidence peaked later in time (∼300 ms) and more centrally.

Selective attention has been hypothesized to be an important driving mechanism, for selective integration models (Glickman et al., 2018) and related theoretical accounts (Bhatia, 2013; Gluth et al., 2018). Our results seem to suggest an important contribution of selective attention in multiple ways. First, contralateral alpha power suppression has long been related to visuospatial attention towards a target (Sauseng et al., 2005; Bacigalupo and Luck, 2019). We found that 375 ms after each stimulus onset there was a suppression of alpha power contralateral to the winning sample. Although later in time than the modulation we found in broadband signals, previous studies have found similar ‘late’ suppression of alpha. Second, the timing, location and pattern of the parametric modulation resemble the findings of a previous study that measured fluctuations in human perceptual choice under focused and divided attention (Wyart et al., 2015). In that experiment, participants integrated tilt information from two simultaneous streams of Gabor patches under focussed attention (where the decision-relevant stream was signalled in advance) and divided attention (where it was not). In the focused condition, posterior electrodes contralateral to the attended stream, but not the unattended stream, showed a negative modulation around 250 ms. The reduced encoding of the losing sample in our study could similarly point to a focussing of attention towards the winning sample. Here we additionally show that this putative neural correlate of attention can be shifted rapidly between the two sides of the screen as samples arrive. Relatedly, the pattern of a posterior negativity contralateral to the target (in this case winning sample) shows a remarkable resemblance to the N2pc component in ERP literature. This component has long been related to (covert) attention and shifts in attention (Luck and Hillyard, 1994; Luck, 2012). Finally, when assessing how eye movements were influenced by decision information, we found that the position of the winning sample did shift the eyes starting from around 300 ms, despite our control for (larger) eye movements. This suggests a possible role of overt attention in selective integration alongside covert attention, although it is worth noting that eye movements only occurred after the observed posterior modulation of neural signals.

While our neural findings support selective integration as described by Tsetsos et al. (2012), they are also in line with other models making similar theoretical claims. For example, the Associations and Accumulation Model (AAM) (Bhatia, 2013) similarly predicts that attention should be drawn to options where the attribute value is high, which in turn asymmetrically drives choices. The SI model also shares a lot of features with the Multi-alternative Decision by Sampling (MDbS) model (Noguchi and Stewart, 2018). One characterizing feature of the MDbS model is that attribute values are integrated through ordinal pairwise comparison, simply tallying attribute ranks. This is in contrast to the SI model, where the magnitude of this difference influences the strength of the evidence weighting. Both our model fits and the neural results suggest that a comparison between attribute values is not solely a process of ordinal comparison. We did find a (small) additive effect by rank (winner vs loser), encoded both in broadband and time-frequency signals. However, there was a much stronger parametric modulation between winning and losing samples that depended on the magnitude difference of the samples, which supports an additional multiplicative step not posited by MDbS.

In conclusion, our study provides a first step in understanding the neural mechanisms which accompany parallel integration and comparison of discrete samples of information.

## Acknowledgements

This research was funded by a British Academy/Leverhulme Small Research Grant (grant number SG141565). FL is funded by the Clarendon Fund, Department of Experimental Psychology and New College Graduate Studentship. CS is funded by a European Research Council Consolidator Award. KT is funded by a European Research Council Starting Grant. The authors thank Nick Yeung for providing access to the EEG equipment and Keno Jüchems for useful feedback on earlier versions of the manuscript.

